# mRNA processing in mutant zebrafish lines generated by chemical and CRISPR-mediated mutagenesis produces potentially functional transcripts

**DOI:** 10.1101/154856

**Authors:** Jennifer L Anderson, Timothy S Mulligan, Meng-Chieh Shen, Hui Wang, Catherine M Scahill, Shao J Du, Elisabeth M Busch-Nentwich, Steven A Farber

**Affiliations:** Carnegie Institution for Science, Department of Embryology, Baltimore, Maryland, United States of America; University of Maryland School of Medicine, Department of Biochemistry and Molecular Biology, Baltimore, Maryland, United States of America; College of Animal Science and Technology, Shandong Agricultural University, China.; Wellcome Trust Sanger Institute, Wellcome Trust Genome Campus, Hinxton CB10 1SA, United Kingdom; Department of Medicine, University of Cambridge, Cambridge, UK

## Abstract

As model organism-based research shifts from forward to reverse genetics approaches, largely due to the ease of genome editing technology, allow frequency of abnormal phenotypes is being observed in lines with mutations predicted to lead to deleterious effects on the encoded protein. In zebrafish, this low frequency is in part explained by compensation by genes of redundant or similar function, often resulting from the additional round of teleost-specific whole genome duplication within vertebrates. Here we offer additional explanations for the low frequency of mutant phenotypes. We analyzed mRNA processing in seven zebrafish lines with mutations expected to disrupt gene function, generated by CRISPR/Cas9 or ENU mutagenesis methods. Five of the seven lines showed evidence of genomic compensation by means of altered mRNA processing: one through a skipped exon that did not lead to a frame shift, one through nonsense-associated splicing that did not lead to a frame shift, and three through the use of cryptic splice sites. These results highlight the need for a methodical analysis of the mRNA produced in mutant lines before making conclusions or embarking on studies that assume loss of function as a result of a given genomic change. Furthermore, recognition of the types of genomic adaptations that can occur may inform the strategies of mutant generation.

**Author summary:** The recent rise of reverse genetic, gene targeting methods has allowed researchers to readily generate mutations in any gene of interest with relative ease. Should these mutations have the predicted effect on the mRNA and encoded protein, we would expect many more abnormal phenotypes than are typically being seen in reverse genetic screens. Here we set out to explore some of the reasons for this discrepancy by studying seven separate mutations in zebrafish. We present evidence that thorough cDNA sequence analysis is a key step in assessing the likelihood that a given mutation will produce hypomorphic or null alleles. This study reveals that alternative mRNA processing in the mutant background often produces transcripts that escape nonsense-mediated decay, thereby potentially preserving gene function. By understanding the ways that cells avoid the deleterious consequences of mutations, researchers can better design reverse genetic strategies to increase the likelihood of gene disruption.

## Introduction

The recent increased use of reverse genetic approaches has been largely driven by the ease, affordability of construction, and implementation of the CRISPR/Cas9 and TALEN systems. Recent communications recount numerous cases of generated mutations in genes of interest lacking an expected effect on phenotypes [1, 2]. The shift from antisense-based knockdown (morpholinos, RNAi) to mutant generation (gene targeting/TILLING methods) resulted in discrepancies in phenotypes, leading researchers to question the specificity and mechanisms of anti-sense technologies and also the methods by which mutants are generated [3]. A screen for essential genes performed in a human cultured cell line found little correlation between genes identified with short hairpin RNA (shRNA) silencing and CRISPR/Cas9 methods [4]. While genome editing methods, such as the CRISPR/Cas9 and TALENs systems, have proven to be an efficient and effective way to reduce or eliminate gene function, a frequent lack of a mutant phenotype is observed, often explained by genetic compensation. This is a process wherein related genes or pathway members are differentially regulated in the mutants to compensate for the targeted loss of a specific gene [3].

In addition to genetic compensation, other mechanisms to recover the function of genes harboring homozygous mutations involve alternative processing of mRNA (what we refer to here as genomic compensation). For example, variations in essential splice sites (ESS) often lead to loss of function resulting in human disease [5, 6]; however, there are several well described ways that function may be recovered [7-9]. In canonical pre-mRNA splicing, joining exons for a functional product requires the presence of a 5’ splice donor sequence (intronic GU), a branchpoint adenosine, the polypyrimidine tract, and a splice acceptor sequence (intronic AG). Base variations in the ESSs lead to one of four outcomes, in order of frequency: 1) exon skipping, 2) activation of cryptic splice sites, 3) activation of cryptic start sites producing a pseudo-exon within the intron or 4) intron inclusion, in the case of short or terminal introns [10]. Mutations in ESSs that lead to skipped exons may result in transcripts that escape nonsense-mediated decay (NMD), the surveillance system that reduces errors in gene expression, if the exon skip does not lead to a frame shift and premature translation termination codon (PTC) [11]. Cryptic splice sites are present throughout the genome both by chance and through evolution of introns [12] and their activation and use by splicing machinery is typical when exon definitions (such as the natural splice sequences) have been altered [13, 14]. Depending on the location of the cryptic splice site used, and the impact on the sequence and frame, functional transcripts may still be generated.

Nonsense-associated alternative splicing, in which a PTC-containing exon is skipped, may also restore the reading frame of a mutated gene [8]. Again, if the exon skip does not lead to a frame shift and new PTC, and the skipped exon does not contain essential motifs, functional transcripts may be generated. Location of the PTC also determines whether the nascent transcripts will be subject to NMD [15]; however, even though these transcripts escape the surveillance system that detects PTCs, these transcripts may either be functional or aberrant. Translation of the transcripts may result in wildtype or deleterious function.

Recently, discussions on how to produce successful knockout models have been renewed [16-18]. To better inform the generation of future mutations, this report analyzes the genetic consequences of several chemical- and CRISPR-induced zebrafish mutant lines in depth. Zebrafish are amenable to current genome editing methods [19, 20] and are a well-established vertebrate model routinely used to assign functions to genes through the use of classical genetic approaches [21]. To begin to investigate the type and frequency of adaptations that may restore gene function in the context of mutation, we carried out studies of mutant lines that included measuring transcript levels and analyzing mRNA splicing (cDNA sequence) and genomic sequence for the presence and use of cryptic splice or start sites. Of the seven mutant lines presented in this study, we show five examples of potential genomic adaptation to mutations that could result in restoration of functional protein production through altered mRNA processing. Our findings emphasize the need to analyze putative mutant lines for genomic compensation at the level of the mRNA sequence and not assume that a mutation will have the predicted effect on mRNA and/or loss of function.

## Results

To investigate how a functional product could be made from a gene containing a putative loss-of-function mutation, we randomly selected seven mutant zebrafish lines generated through chemical- or CRISPR-mediated mutagenesis to study (Table 1). Six zebrafish lines carrying mutations in genes involved in lipid metabolism were obtained through the Zebrafish Mutation Project (ZMP, Wellcome Trust Sanger Institute) (*abca1a^sa9624^, abca1b^sa18382^*, *cd36*^sa9388^, *creb3l3a^sa18218^, pla2g12b^sa659^*, and *slc27a2a^sa30701^).* Lines were generated with point mutations throughout the genome using classical ENU mutagenesis [22], followed by association of the induced mutations with protein-coding genes using whole exome sequencing methods [1]. We selected five lines that had mutations in essential splice sites and one line with a nonsense mutation (*creb3l3a^sa18218^)*, with the aim to investigate the genome’s ability to compensate for induced mutations through the generation of novel alternative transcripts. In addition to the 6 ZMP lines with ENU-induced base changes, a line with a CRISPR/Cas9-generated deletion was included. cDNA sequence and transcript levels of pooled wildtype and homozygous mutant larvae were analyzed.

**Table 1.** Mutations in this study.

| Gene mutated | Ensembl ID | Allele | Nature of mutation | Method of mutation |
| --- | --- | --- | --- | --- |
| *abca1b* | ENSDARG00000079009 | *sa18382* | point mutation in ESS | ENU |
| *slc27a2a* | ENSDARG00000036237 | *sa30701* | point mutation in ESS | ENU |
| *abca1a* | ENSDARG00000074635 | *sa9624* | point mutation in ESS | ENU |
| *cd36* | ENSG00000135218 | *sa9388* | point mutation in ESS | ENU |
| *pla2g12b* | ENSG00000138308 | *sa659* | point mutation in ESS | ENU |
| *creb3l3a* | ENSDARG00000056226 | *sa18218* | nonsense mutation | ENU |
| *smyd1a* | ENSDARG00000009280 | *mb4* | 7-bp deletion in exon | CRISPR/Cas9 |

### Two of the five ZMP lines with ESS mutations lose the predicted exon

To determine whether each ZMP line containing an ESS mutation results in the predicted skipping of an exon, adult heterozygote mutant zebrafish were incrossed and their offspring were pooled or individually processed into total RNA and cDNA. PCR amplification of this cDNA template revealed amplicons of the expected size from each ZMP line, while *abca1b^sa18382^* (ATP-binding cassette transporter, sub-family A, member 1B) and *slc27a2a^sa30701^* (solute carrier family 27, member 2a; protein is fatty acid transport protein, member 2a) also had a shorter amplicon (116 and 210 bp shorter, respectively; S1 Fig) that matches the predicted length of amplicons of these cDNAs after the omission of the affected exon. The *pla2g12b^sa659^* (phospholipase A2, group XIIB) mutant allele has a mutation in the essential splice acceptor site preceding its final (fourth) exon and could not be investigated for the loss of that exon using these methods.

To confirm that exons are skipped in *abca1b^sa18382^* and *slc27a2a^sa30701^* and determine why the mutations in *abca1a^sa9624^* (ATP-binding cassette transporter, sub-family A member 1A) and *cd36^sa9388^* (cluster of differentiation 36, aka fatty acid translocase) did not appear to lead to the predicted skipping of exons (S2 Fig), individual 6-dpf larvae underwent genotyping, gDNA and cDNA sequence analysis, and qPCR studies.

### An ESS mutation in *abca1b* leads to a skipped exon and early termination signal

*abca1b^sa18382^* has a point mutation in the essential splice acceptor site of intron 33–34 (g.64427G>T) (Fig 1). To determine whether the point mutation results in skipping of the subsequent exon (e34) and use of the (next) essential splice acceptor site of intron 34–35, we performed PCR amplification using primers targeted to flanking exons of cDNA (synthesized from individual, genotyped larvae; 713-bp amplicon), followed by Sanger sequencing. As expected, cDNA sequencing confirmed that exon 34 (116 bases) is skipped and leads to a frame shift in *abca1b*^sa18382/+^and *abca1b^sa18382/sa18382^* larvae. Following the frame shift, the mutant cDNA encodes a 13 AA open reading frame (ORF) and an early termination signal that would direct the loss of exons 34–46. qPCR studies reveal transcript levels are down 3.5-fold in 6-dpf *abca1b^sa18382/sa18382^* zebrafish (ANOVA with Tukey’s test, p=0.049).

**Fig 1.**
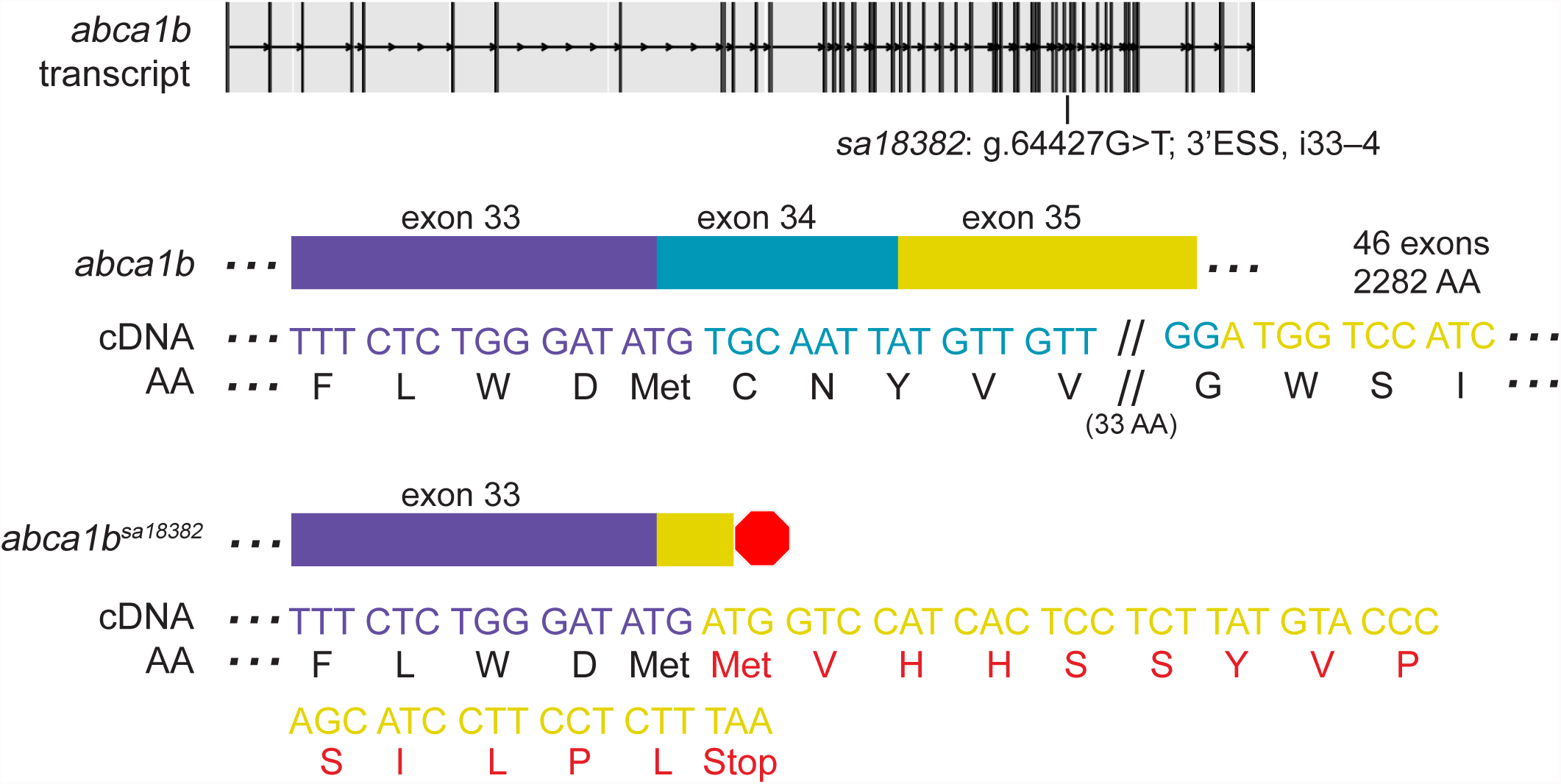
An ENU-induced G>T mutation in the 3’ essential splice site (ESS) of i33–34 of *abca1b* leads to a skipped exon and an early termination signal. Analysis of cDNA sequence and agarose gel electrophoresis (Supplemental Figure 1) indicates that loss of exon 34 (116 bases) causes a frame shift, followed by a short ORF (13 AA) and an early termination codon (shown in red). qPCR studies revealed transcript levels down 3.5-fold in 6-dpf *abca1b^sa18382/sa18382^* zebrafish. See Supplemental Table 1 for sequence spanning mutation and predicted outcome.

### An ESS mutation in *slc27a2a* leads to an expected skipped exon but not a frame shift

*slc27a2a^sa30701^* has a point mutation in the essential splice donor site of intron 2– 3 (g.3431G>A) (Fig 2). cDNA sequencing confirms omission of exon 2 in *slc27a2a*^sa30701/+^and *slc27a2a^sa30701/sa30701^* larvae. No frame shift is observed since exon 2 is 210 bases long (encoding 70 AA). By qPCR, transcript levels in 6-dpf *slc27a2a^sa30701/sa30701^* zebrafish did not differ from those of their wildtype siblings (ANOVA with Tukey’s test; p-value greater than threshold of 0.05).

**Fig 2.**
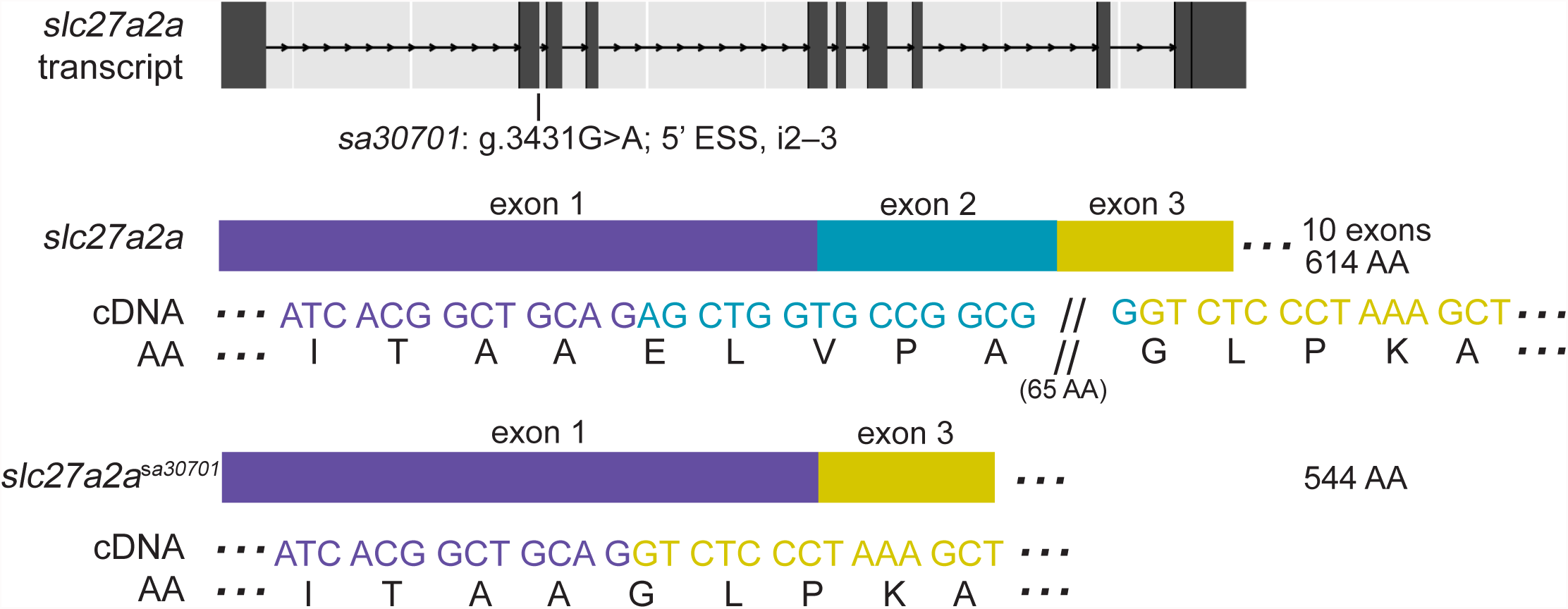
An ENU-induced G>A mutation in the 5’ essential splice site (ESS) of i2–3 of *slc27a2a* leads to a skipped exon. Analysis of cDNA sequence and agarose gel electrophoresis (Supplemental Figure 1) reveals a loss of exon 2 (70 AA) with no frame shift. By qPCR, transcript levels of 6-dpf *slc27a2a^sa30701/sa30701^* zebrafish were not found to be different than their siblings. See Supplemental Table 1 for sequence spanning mutation and predicted outcome.

### An ESS mutation in *abca1a* leads to use of a nearby cryptic splice acceptor site and loss of a single non-essential AA

*abca1a^sa9624^* has a point mutation in the 3’ ESS of intron 29–30 (g.48320G>A) (Fig 3). Analysis of cDNA sequence from individual genotyped larvae revealed the loss of three bases, “TAG”, at the start of exon 30 in heterozygous and homozygous mutants. To look for cryptic splice sites, a flanking region of gDNA was PCR amplified and sequenced. A cryptic splice acceptor site (“AG”) was found 2 and 3 bases downstream of the mutated wildtype splice acceptor site, in exon 30. Use of this cryptic splice acceptor site splices out the first three bases of exon 3 (“TAG”) and the protein this message encodes would lack one Serine (and remain in frame with the wildtype product). Transcript levels in 6-dpf *abca1a^sa9624/sa9624^* zebrafish did not differ from their wildtype siblings (ANOVA with Tukey’s test; p-value>0.05).

**Fig 3.**
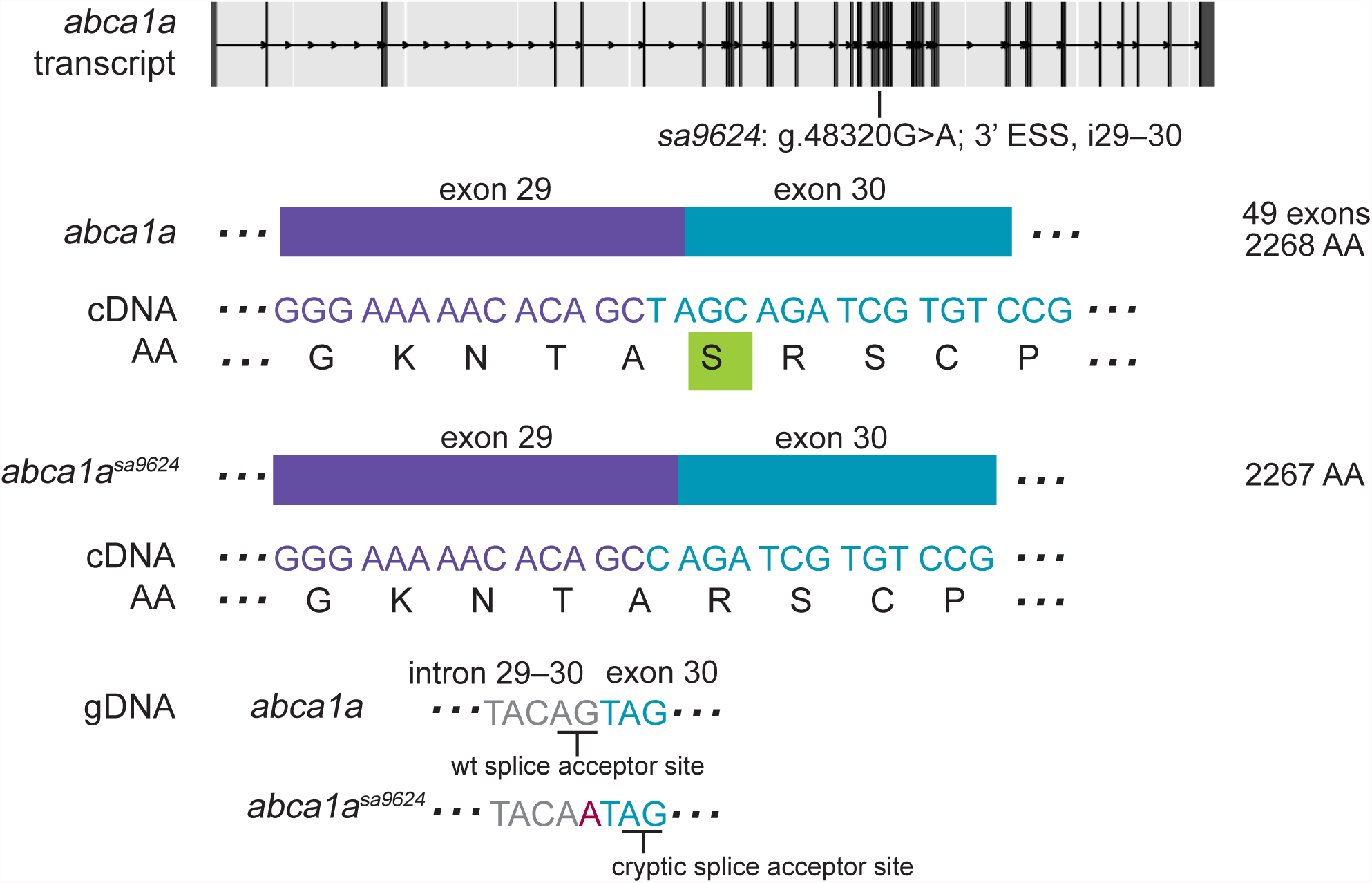
An ENU-induced G>A mutation in the 3’ essential splice site (ESS) of i29–30 of *abca1a* leads to use of a nearby cryptic splice site and loss of a single AA. The base change causes a missed splice acceptor and sequence immediately following the mutated base to be used as a cryptic splice site, as confirmed by analysis of cDNA sequence. A single Serine is lost (boxed in green) and the product remains in frame. By qPCR, transcript levels of 6-dpf *abca1a^sa9624/sa9624^* zebrafish were not found to be different than their siblings. See Supplemental Figure 2 for agarose gel electrophoresis of amplified cDNA and Supplemental Table 1 for sequence spanning mutation and predicted outcome.

### An ESS mutation in *cd36* leads to use of a cryptic splice donor site, frame shift, PTC, but not NMD

*cd36^sa9388^* has a point mutation in the 5’ ESS (splice donor site) of intron 10–11 (g.11242G>A) (Fig 4). cDNA sequencing of individual, genotyped larvae reveals incorporation of extra bases “ATAT” in between the sequence for exon 10 and exon 11, which leads to a frame shift in the mutant allele. After the frame shift, 18 AA and a PTC follow, predicting the loss of exon 12 (154 AA). The PTC position sits at the last exon-exon junction and thus transcripts are predicted to escape NMD. Transcript levels of 6-dpf *^cd36sa9388^*^/*sa9388*^ larvae did not differ significantly from their wildtype siblings (ANOVA with Tukey’s test; p-value>0.05).

**Fig 4.**
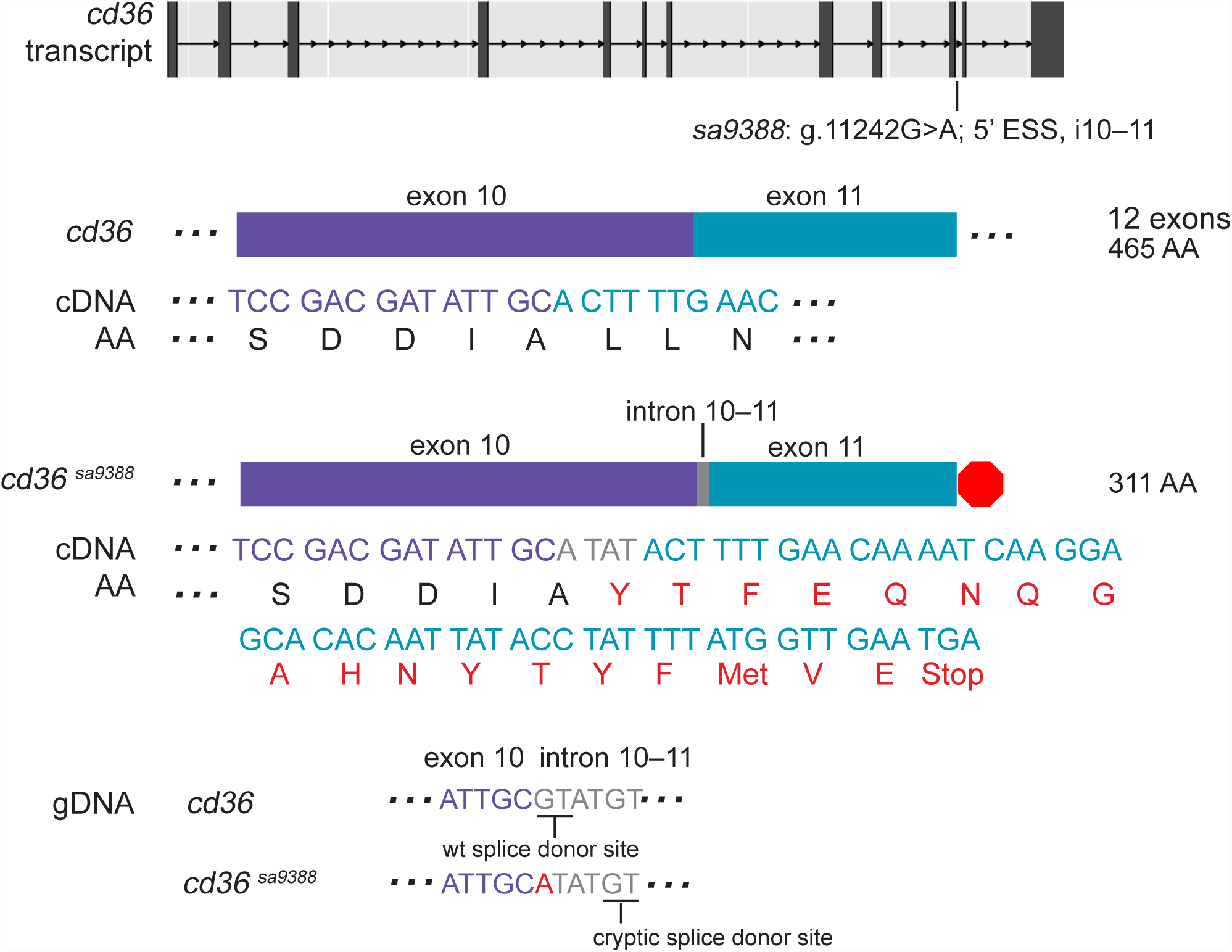
An ENU-induced G>A mutation in the 5’ essential splice site (ESS) of i10–11 of *cd36* leads to use of a cryptic splice site and a frame shift. The base change causes loss of a splice donor and use of a cryptic splice site 3 and 4 bases downstream, as confirmed by analysis of cDNA sequence. The intronic sequence “ATAT” preceding the cryptic splice site is thus incorporated, leading to a frame shift. After the frame shift, 18 AA follow before a stop codon (shown in red) directs early termination and loss of exon 12 (154 AA). By qPCR, transcript levels of 6-dpf *cd36sa9388/sa9388* zebrafish were not found to be different than their siblings. See Supplemental Figure 2 for agarose gel electrophoresis of amplified cDNA and Supplemental Table 1 for sequence spanning mutation and predicted outcome.

To look for the use of a cryptic splice donor site, a flanking region of gDNA isolated from individual larvae was amplified and sequenced. The wildtype sequence at the 5’ end of intron 10–11 includes the splice donor “GT”. However, in the mutant allele, the first base is mutated to an “A”, resulting in the loss of the splice donor site. The mutated intronic sequence begins “ATATGT…”, which provides a cryptic splice donor site (“GT”) 3 and 4 bases downstream of the mutated wildtype splice donor site (Fig 4).

### An ESS mutation in *pla2g12b* is associated with lowered transcript counts and a mutant phenotype

*pla2g12b^sa659^* has a point mutation in the splice acceptor site of intron 3–4 (g.10194A>T) and is predicted to skip the last (4 of 4) exon; thus, exons flanking the mutation could not be PCR amplified to confirm the loss of exon 4 in mutants. Attempts to amplify an alternative transcript with retention of either the final exon 4 or the intron 3–4 did not succeed when using cDNA synthesized from homozygous mutant larvae as the template. During phenotypic screening and genotyping of 5-dpf larvae from heterozygous incrosses, a total of 29 *pla2g12b^sa659/sa659^* larvae exhibited a darkened yolk phenotype while 52 *pla2g12b^+/+^* siblings did not (2 experiments) (phenotype: http://www.sanger.ac.uk/sanger/ZebrafishZmpgene/ENSDARG00000015662#sa659). Correspondingly, RNA expression profiling methods reflect a 3-fold decrease in mutants/siblings (p=3.96 x 10^−6^) as determined using the method of Anders and Huber (2010) [23] (expression profile from: http://www.sanger.ac.uk/sanger/ZebrafishZmpmRNAexpression/45).

## A nonsense mutation in *creb3l3a* leads to an unexpected skipped exon but no frame shift

*creb3l3a^sa18218^* has a nonsense mutation in exon 2 of 10 (g.357C>T), which changes codon <u>C</u>AA to <u>T</u>AA, a PTC (Fig 5). PCR amplification of cDNA (wildtype and homozygous pooled larval intestines) followed by gel electrophoresis revealed two bands in homozygous mutants but only the expected wildtype band in the wildtype siblings (Fig 5). cDNA sequencing of the bands showed alternative transcripts with the unexpected omission of exon 2 in homozygous mutant but not in wildtype larvae. Splicing out exon 2 (114bp encoding 38 AA) does not lead to a frame shift. The nonsense mutation was found to occur in a predicted exonic splice enhancer (ESE) sequence using the web-based prediction tool, ESEFinder [24]. Mutation of an ESE, an important aspect of exon definition, could explain the failure of the mutated cDNA to include exon 2. By qPCR, transcript levels of 6-dpf *creb3l3a^sa18218/sa18218^* dissected intestines did not differ significantly from their wildtype siblings (Wilcoxon-Mann-Whitney test; p>0.05).

**Fig 5.**
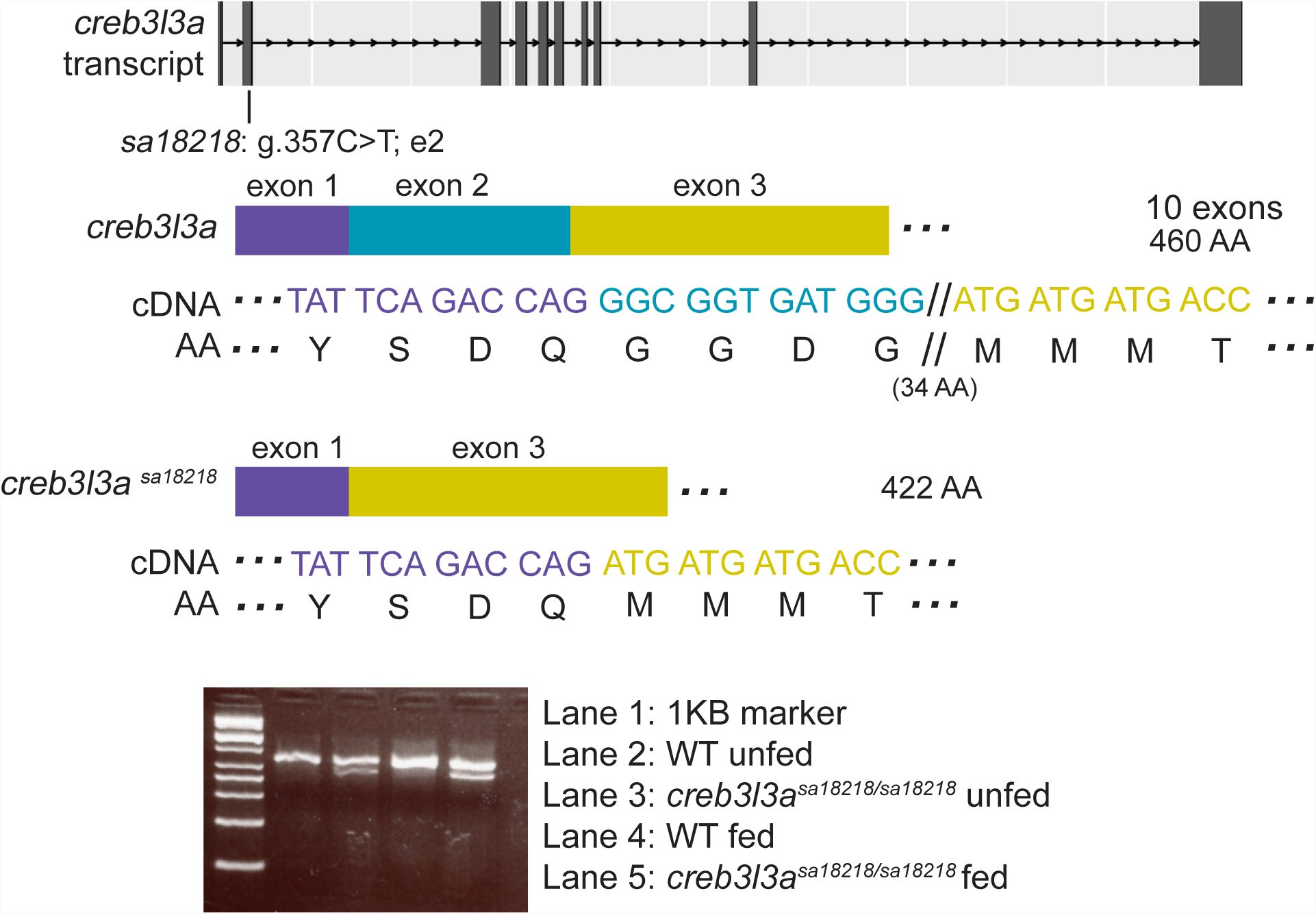
An ENU-induced C>T nonsense mutation in exon 2 of *creb3l3a* leads to a skipped exon. Analysis of cDNA sequence reveals a loss of exon 2 (38 AA) with no frame shift. Agarose gel electrophoresis of amplified cDNA (from pooled intestines, fed and unfed) revealed an additional band in the homozygous mutants matching the expected size of a product with a skipped exon 2. By qPCR, transcript levels of 6-dpf *creb3l3a^sa18218/sa18218^* zebrafish were not found to be different than their siblings. See Supplemental Table 1 for sequence spanning mutation and predicted outcome.

### A CRISPR-induced deletion in *smyd1a* correlates with the use of upstream cryptic splice sites

To confirm that genomic adaptation is not a phenomenon specific to ENU-mutagenized lines, we analyzed a 7-bp deletion in exon 3 of *smyd1a* (g.6948_6955del; SET and MYND domain containing 1A) which was generated using CRISPR/Cas9 targeting methods. The 7-bp deletion leads to a predicted frame shift and PTC (48/485 AA produced) (Fig 6). However, *smyd1a^mb4/mb4^* larvae did not display a mutant phenotype as determined by whole-mount immunostaining at 28 hpf using the F59 antibody which recognizes slow myosin heavy chain specifically expressed in zebrafish slow myofibers; myosin expression and sarcomere organization were the same in the mutants and wildtype controls. To look for evidence of novel alternative splicing, *smyd1a* cDNA was sequenced from wildtype and mutant embryos by cloning full-length PCR products. As expected, all 20 wildtype cDNA clones had the *smyd1a* wildtype sequence and all 20 clones from the homozygous mutant embryos contained the 7-bp deletion in exon 3. However, 6 of the 20 cDNA clones from mutant embryos exhibited alternative splicing at exon 2 (Fig 6). Three clones had an alternative splice event using a cryptic splice acceptor site (“AG”) in exon 2, located 13-bp downstream of the wildtype splice acceptor site, leading to a 13-bp deletion at the 5’ end of the exon 2. Similarly, sequence data from another three clones show the use of a cryptic splice acceptor site (“AG”) 40 bp downstream of the wildtype splicing site, resulting in a 40-bp deletion at the 5’ region of exon 2. Both deletions are predicted to lead to a frame shift and premature translation termination. qPCR studies revealed transcript levels of 1- and 2- dpf *smyd1^amb4/mb4^* zebrafish were down 13-fold compared to wildtype siblings (Wilcoxon-Mann-Whitney test, p=0.000077).

**Fig 6.**
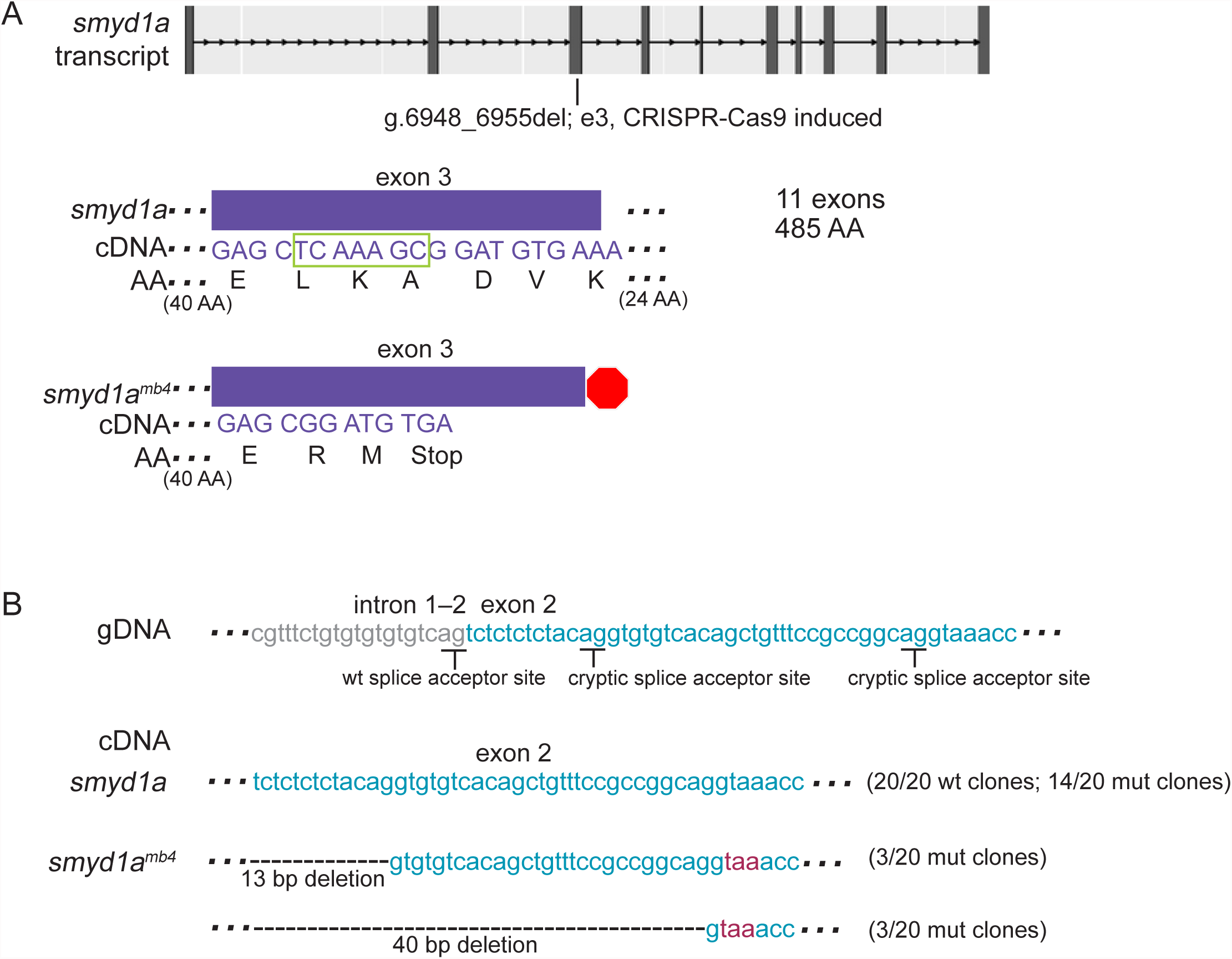
A CRISPR-induced deletion (7 bp) in exon 3 of *smyd1a* corresponds to use of cryptic splice sites in exon 2 in mutant clones. **A.** A 7-bp deletion in exon 3 (boxed in green) is predicted to lead to a frame shift mutation and early termination. Of 20 mutant clones sequenced, all 20 had the expected 7-bp deletion. **B.** In addition, 3 revealed use of one cryptic splice site and another 3 revealed use of a second cryptic splice site in exon 2, leading to a 13- and 40-bp deletion, respectively. Both deletions result in a frame shift and premature termination codon, shown in red text. 20 randomly selected wt clones did not show alternative splicing. By qPCR, transcript levels of 1- and 2-dpf *smyd1^amb4/mb4^* zebrafish were down thirteen-fold compared to wildtype siblings. See Supplemental Table 1 for sequence spanning mutation and predicted outcome.

## Discussion

In this report, we analyzed the genomic adaptations that arise in zebrafish in response to ENU- and CRISPR-induced mutations (Table 2). Recently, Popp et al. reviewed how the process of exon-junction-complex-mediated NMD influences the success of creating loss-of-function mutations with CRISPR/Cas9 [17]. Most notable is their earlier finding that NMD cannot occur if a PTC is within 50–55 nucleotides (nt) of the last exon-exon junction [15]. In our study, we found one example of this phenomenon. For the *cd36^sa9388^* allele, the resultant PTC is within 1 nt of the last exon junction (e11–e12) and as predicted, we observed wildtype transcript levels in the homozygous mutants. Another group has proposed identifying potential cryptic start sites before the construction of any CRISPR or TALENs vectors, after finding wildtype expression levels in six separate *in vitro* mouse NIH3T3 cell lines harboring frame-shift mutations in Gli3 [16]. Loss of function from mutations near the translation initiation site may be recovered by utilizing nearby downstream alternative translation initiation sites [25]. The mutations in our lines were closer to the middle or 3’ end of genes. We did identify use of alternative splice sites in the mutant allele in three of seven lines (*abca1a^sa9624^*, *cd36^sa9388^, smyd1a^mb4^*), underlining the importance of identifying potential cryptic splice sites prior to basing studies on presumed lack of gene function. While alternative splicing in the *smyd1a^mb4^* line is predicted to lead to a PTC, we do not observe a mutant phenotype. Lack of phenotype in *smyd1a^mb4/mb4^* larvae suggests transcripts may retain function or that related genes or pathway members compensate for the loss of function. We also found an example of nonsense-associated splicing (*creb3l3a^sa18218^*), wherein a PTC-containing exon is spliced out and *creb3l3a^sa18218/sa18218^* larvae have wildtype transcript levels. The mechanisms underlying this process are still being explored: in many cases, mutations in conserved splice elements (such as exonic splice enhancers; ESE) have been shown to cause nonsense-associated splicing [26-30]. Prykhozhij et al. also recently illustrated the need for careful mutation analysis, beyond the level of gDNA sequence. They found only one of three mutant zebrafish lines resulted in the predicted frameshift [18, 31]. Of the remaining two lines, one displayed an exon skip, possibly due to a mutation in an ESE, and the other used an alternative start site.

**Table 2.** Summary of outcomes from our study.

| Gene mutated | Ensembl ID | Allele | Nature of mutation | Method of mutation | Outcome on cDNA sequence | Outcome on mutant transcript levels |
| --- | --- | --- | --- | --- | --- | --- |
| *abca1b* | ENSDARG00000079009 | *sa18382* | point mutation in ESS | ENU | Skipped exon (116 bp), frame shift, PTC | 3.5-fold down |
| *slc27a2a* | ENSDARG00000036237 | *sa30701* | point mutation in ESS | ENU | Skipped exon (210 bp), frame maintained | WT levels |
| *abca1a* | ENSDARG00000074635 | *sa9624* | point mutation in ESS | ENU | Downstream cryptic splice site used, loss of single Serine, frame maintained | WT levels |
| *cd36* | ENSG00000135218 | *sa9388* | point mutation in ESS | ENU | Downstream cryptic splice site used, frame shift, PTC | WT levels |
| *pla2g12b* | ENSG00000138308 | *sa659* | point mutation in ESS | ENU | No evidence found for retention of exon or preceding intron | 3-fold down |
| *creb3l3a* | ENSDARG00000056226 | *sa18218* | nonsense mutation | ENU | Skipped affected exon (114 bp), frame maintained | WT levels |
| *smyd1a* | ENSDARG00000009280 | *mb4* | 7-bp deletion in exon | CRISPR/Cas9 | (Deletion) frame shift, PTC. Two cryptic splice sites used in mutant clones, frame shift, PTC | 13-fold down |

Since ESS mutations often lead to human disease [6], *in vivo* models are critical to our understanding. However, we found that skipping an exon may still lead to a viable product: if the exon is divisible by 3 and thus its omission does not lead to a frame shift and PTC, transcript levels were not subject to NMD (*creb3l3a^sa18218^, slc27a2a^sa30701^)*. In both of these lines, sequence alignment with the human orthologue revealed no essential motifs in the skipped exons. Examination of intron-spanning reads from available temporal expression data revealed no evidence of the alternative transcripts we identified in this study in wildtype larvae [32], suggesting that they did not result from wildtype alternative splicing events.

In this study, we report that five of seven analyzed zebrafish lines with induced mutations show evidence of genomic adaptation and contribute to the growing data of how to produce successful knockout models. Our data support a hypothesis that there may be a surveying mechanism that could detect mutations and adapt mRNA alternative splicing to cope with the loss of function. Analysis of cDNA sequence in mutant alleles may allow for prediction of genomic adaptation, simply by scanning for proximal cryptic splice and initiation sites that might be used for alternative transcripts.

Employing multiple “guide” RNAs in the CRISPR/CAS9 system can result in large intron-spanning deletions in or the elimination of targeted genes. While this approach has been used to generate loss-of-function alleles, it can lead to the deletion of the genomic regions needed for post-transcriptional regulation of gene expression or transcriptional regulation of other genes. It is estimated that 30–80% of human coding genes are post-transcriptionally regulated, at least in part, by microRNAs (miRNAs) [33]; so far, 2,619 miRNAs and 324,219 miRNA-target interactions have been annotated in human (miRTarBase) [34] and approximately 50% of miRNA coding sequence is located in the introns of coding genes [35]. Rather than creating large deletions or removing an entire gene, other approaches, such as those used to generate nonsense mutations or small deletions, may work better to generate loss-of-function alleles that retain these regulatory regions. As we have shown, alternative transcripts may escape nonsense-mediated decay so 1) analyze the DNA sequence for nearby cryptic splice sites, especially those in frame to the natural/altered cryptic splice site, 2) check whether a nonsense mutation is in a predicted splicing enhancer sequence using available web tools, and 3) in the case of expected exon skip, analyze the exonic sequence for essential motifs and whether the exon length is divisible by 3. Since shorter introns that precede expected affected exons may be retained (instead of exon skip), intron length is also a factor to consider when generating mutants. Performing these steps near the start of a project can inform the nature and location of mutations that would most likely result in a loss-of-function mutant with a phenotype of interest.

## Materials and Methods

### Zebrafish husbandry

All procedures in this study were approved by the Carnegie Institution Animal Care and Use Committee (Protocol# 139) or the Institutional Animal Care and Use Committee of the University of Maryland (Permit Number: 0610009). All lines were raised and crossed according to zebrafish husbandry guidelines [36].

### Genotyping carriers (ZMP lines)

Heterozygotes for each mutation were identified through a fin-clip based gDNA isolation (REDExtract-N-Amp Tissue PCR kit; Sigma-Aldrich), PCR amplification of a 400–600 bp region around the mutation using designed primer sets (MacVector, Primer 3), and Sanger sequencing using a nested sequencing primer. (Primer sets and conditions are in S2 Table.)

For the creb3l3a^sa18218^ line, an NaOH-based DNA extraction method was used to extract gDNA from fin tissues. Genotyping primers were designed using dCAPS finder 2.0 with one mismatch (http://helix.wustl.edu/dcaps/dcaps.html; [37]). The primer introduces EcoRV restriction sites in the mutant amplicons but not in the WT amplicons.

### To look for evidence of a skipped exon in the lines with mutations in essential splice sites (ZMP lines)

For each line with a mutation in an ESS, larvae were collected from incrosses of identified heterozygotes and 10–20 6-dpf larvae were pooled for generating RNA samples (using above protocol). RNA samples served as template to generate cDNA (iScript cDNA Synthesis Kit, Bio-Rad). cDNA samples were PCR amplified to provide amplicon sizes of 400–700 bp) and the products were separated and sized using gel electrophoresis. For lines that showed evidence of a skipped exon, individual larvae were genotyped and treated similarly to above to correlate amplicon size with genotype.

### To generate cDNA after genotyping individual larvae for qPCR analysis (ZMP lines)

Adults that were found to carry the ESS mutation were incrossed and the progeny collected. Individual 6-dpf larvae underwent a Trizol-based RNA prep adapted from Macedo and Ferreira (2014) [38] to include an additional chloroform extraction. To genotype individual samples, residual gDNA in the unpurified RNA samples was PCR amplified and sent for Sanger sequencing. After genotypes were determined (SnapGene Viewer to view peak trace files), RNA samples were DNAase I-treated and purified (RNA Clean and Concentrator, Zymo Research), served as templates for cDNA synthesis (iScript cDNA Synthesis Kit, Bio-Rad), and ultimately used in qPCR studies to analyze transcript levels.

### To check transcript levels using Real-time PCR

qPCR methods included SYBR Green-based methods (Sigma-Aldrich, *abca1a, creb3l3a, smyd1a*) and Taqman gene expression assays (ThermoFisher Scientific; *cd36*, *slc27a2a*, and *abca1b*). ef1α (for *smyd1a*) or 18s rRNA (for all others) levels were used as reference genes. Primer and assay information is shown in S2 Table.

cDNA samples for individual larvae, along with “No RT” controls and “No transcript” controls were run on the CFX96 Touch Real-Time PCR Detection System (Bio-Rad) or on the 7500 Fast-Time PCR System (AB Applied Biosystem). Three technical replicates were run for each sample and a minimum of three biological replicates were used for each genotype for each line. Data was analyzed through calculation of Delta Ct values (18s rRNA as internal control) and either one-way analysis of variance (ANOVA) with the Tukey post hoc test or the Wilcoxon-Mann-Whitney test [39].

### To perform transcript counts

Transcript counting data for the *pla2g12b* mutant line (sa659) was obtained from the Sanger Zebrafish Mutation Project [1] and performed as described [32]. Statistical significance tests were performed using the method of Anders and Huber (2010) [23].

### To analyze cDNA sequencing (ZMP lines)

cDNA for individual larvae was generated as described above and sequencing was obtained using Sanger sequencing methods. Peak trace files were analyzed manually using SnapGene Viewer (GSL Biotech, LLC) or MacVector. To assist in determining the two alleles of interest (wildtype and potential mutant) for each line, Poly Peak Parser [40] and alignment of wildtype and mutant alleles in MacVector (Align to Reference) were used.

### Generation of the smyd1a^mb4^ mutant

The *smyd1a* allele containing a 7-bp deletion was generated using the CRISPR/Cas9 targeting method (Cai and Du, in preparation). The target site (5’-GGACCTGAAGGAGCTCAAA-3’) was located in exon 3 of the *smyd1a* gene. Genotyping was carried out by using gDNA extracted from the caudal fin as a template for PCR followed by SacI digestion of the resulting amplicons. The 7-bp deletion abolished the SacI site, allowing resolution of bands by agarose gel electrophoresis.

### DNA and mRNA sequence analysis of the smyd1a^mb4^ mutant

Homozygous *smyd1a* mutants were identified by PCR and SacI digestion, and confirmed by sequencing the PCR product. To compare the *smyd1a* mRNA sequences from wild type (WT) and mutant embryos, total RNA was isolated from a pool of 50 WT and homozygous *smyd1a* mutant embryos at 48 hpf. cDNAs were generated using the RevertAid First Strand cDNA Synthesis Kit (ThermoFisher, K1621). The full length *smyd1a* cDNA was amplified from the WT and mutant template using Phusion^®^ High-Fidelity DNA Polymerase (NEB, M0530S). The amplicons were A-tailed using Taq DNA polymerase (Promega, M8295) and subsequently cloned into pGEM-T easy (Promega, A1360).

## Acknowledgments

J.L.A., C.M.S, E.M.B. and S.A.F. were supported in part by the Carnegie, Duke and Sanger Center consortium to identify zebrafish genes required for lipid processing and digestive organ function (Farber PI, Rawls and Busch-Nentwich Co-PIs; National Institute of Diabetes and Digestive and Kidney [NIDDK] R01DK093399) and National Institute of General Medicine (GM) R01GM63904 to the Zebrafish Functional Genomics Consortium (Stephen Ekker PI and S.A.F. Co-PI). The content is solely the responsibility of the authors and does not necessarily represent the official views of the National Institutes of Health (NIH). E.M.B. was funded by the Wellcome Trust Sanger Institute (grant number WT098051). S.D. was supported by a research fund from University of Maryland Baltimore. Shandong Provincial Education Association for International Exchanges provided a visiting professor fellowship to H.W. Additional support for this work was provided by the Carnegie Institution for Science endowment and the G. Harold and Leila Y. Mathers Charitable Foundation to the laboratory of S.A.F. We thank Fred Tan for the software used to generate transcript illustrations.

## Supporting information captions

**S1 Fig. Two lines with ESS mutations have a shorter amplicon, supporting an exon skip.** *slc27a2a^sa30701^* (solute carrier family 27, member 2a) and *abca1b^sa18382^* (ATP-binding cassette transporter, sub-family A, member 1B) had a shorter amplicon (210 and 116 bp shorter, respectively) that matches the predicted length of amplicons of these cDNAs (pooled from heterozygous incrosses) after the omission of the affected exon. A. Red arrows indicate shorter amplicons. B. Primers used and sizes anticipated are listed. C. Primer locations are indicated by arrows. The exons that are expected to be skipped as a result of ESS mutations are shown in yellow.

**S2 Fig. Three lines with ESS mutations do not reveal shorter amplicons expected with an exon skip.** *abca1a^sa9624^* (ATP-binding cassette, sub-family A, member 1A), *cd36^sa30701^* (cluster of differentiation cd), and *pla2g12b^sa659^* (phospholipase A2 Group XIIB) yield PCR products (using pooled larvae from heterozygous incrosses) that match the predicted length of amplicons of the wildtype cDNAs and do not reveal a shorter amplicon expected with the omission of the affected exon (A). Modifying PCR conditions to look for evidence of a retained intron (i3–4) in *pla2g12b^sa659^*did not yield a product in homozygous mutants. B. Primers used and sizes anticipated are listed. C. Primer locations are indicated by arrowheads or arrows. The exons that are expected to be skipped as a result of ESS mutations are shown in yellow. For *pla2g12b^sa659^*, the splice acceptor sequence preceding exon 4 (shown in gray) is mutated (and there is no downstream natural splice acceptor sequence).

**S1 Table.** Mutations analyzed in this study and their predicted outcome.

**S2 Table.** List of primers and methods used in this study.

